# Membrane composition and lipid to protein ratio modulate amyloid kinetics of yeast prion protein

**DOI:** 10.1101/2020.08.07.241299

**Authors:** Arnab Bandyopadhyay, Achinta Sannigrahi, Krishnananda Chattopadhyay

## Abstract

Understanding of prion aggregation in membrane environment may help to ameliorate neurodegenerative complications caused by the amyloid forms of prions. Here, we investigated the membrane binding induced aggregation of yeast prion protein Sup35. Using the combination of fluorescence correlation spectroscopy (FCS) at single molecule resolution and other biophysical studies, we establish that lipid composition and lipid/protein ratio are key modulators of the aggregation kinetics of Sup35. In the presence of zwitterionic membrane, Sup35 exhibited a novel biphasic aggregation kinetics at lipid/protein ratio ranging between 20:1 and 70:1 (termed here as the Optimum Lipid Concentration, OLC). In ratios below (Low Lipid Concentration, LLC) and above (ELC, Excess Lipid Concentration) that range, the aggregation was found to be monophasic. In contrast, in the presence of negatively charged membrane, we did not observe any bi-phasic aggregation kinetics in the entire range of protein to lipid ratios. The toxicity of the aggregates formed within OLC range was found to be greater. Our results provide a mechanistic description of the role that membrane-concentration/composition-modulated-aggregation may play in neurodegenerative diseases.

## Introduction

Prion proteins are involved in several fatal and transmissible neurodegenerative diseases, including transmissible spongiform encephalopathies, human Creutzfeldt-Jacob disease and sheep scrapie, in which they have been shown to transmit their misfolded shape onto normal variants ^*1, 2*^.On microscopic examination, the brains of patients with prion disease typically reveal characteristic histopathologic changes, consisting of neuronal vacuolation and degeneration^*3*^. In humans, a conformational change of the prion protein between its non-toxic PrP^c^ and a toxic PrP^sc^ form is believed to be responsible for the disease pathology^*4, 5*^. PrP^sc^ contains significant beta-sheet, which facilitates the formation of highly insoluble aggregates. These aggregates are partially resistant to proteolytic digestion^*6*-9^. It is known that amyloidogenic proteins can populate numerous structurally distinguished amyloid forms or strains with distinct phenotypes^*10*^. However, little is known about the structural morphology of the diverse amyloid conformers formed under different environmental conditions.

Because of the presence of glycosylphosphatidylinositol (GPI)-anchor ^*9, 11-13*^, PrP^c^ can be associated with lipid membranes. Whatley et al have shown that the vacuolation in nerve cells is drastically increased during prion aggregation, which further emphasizes the effect of membrane environments towards the aggregation of prion protein^*14*^. Since it is difficult to generate prion strains from solely pure protein de-novo that infect wild type animals and cause transmissible disease, the effect of membrane on distinct infectious PrP amyloid conformers remains unclear. As a result, it is not understood if and how membrane environment would modulate the folding and amyloid formation of the prion protein. Fei wang et al ^*9*^, have found earlier that the PrP-lipid interaction can be initiated by electrostatics, which can be followed by hydrophobic interactions. They show that the protein-lipid interaction can convert full-length R-helix-rich PrP to a PrP^Sc^-like conformation under physiological conditions, supporting the relevance of lipid-induced PrP conformational change to in-vivo PrP conversion.

Prion like proteins are also found in different fungi, including yeast^*15*^. Sup35 is the yeast version of the eukaryotic translation factor 3 (eRF-3), which is involved in the termination of translation in Saccharomyces cerevisiae^*16, 17*^. Sup35 is not infectious to humans, although it carries the same transmissible phenotype as the human prion protein^*18*^. Since there is obvious resemblance between its proposed mechanism of propagation and that of the mammalian prion disease, Sup35 is often used as a model protein to study the prion disease^*19, 20*^. It has been shown that Sup35 may propagate as a prion in mammalian cells and that GPI anchoring facilitates aggregate propagation between N2a cells, resembling mammalian prion behavior ^*21*-24^.

Here, we report a detailed study on the aggregation kinetics of Sup35 protein in the presence of model phospholipid vesicles using FCS and other biophysical methods. FCS offers a sensitive and convenient method to study biomolecular diffusion under single molecule resolution. In this study, we chose a physiologically relevant lipid PC (phosphatidylcholine) and PS (phosphatidylserine) since these are the most abundant components of the membranes of synaptic vesicles ^*25*^. We chose the phospholipid DMPC (14:0 saturated, transition temperature ∼24 °C), since it exists in the fluid phase at physiological relevant temperature of 37°C ^*26*^.The molecular models of cell membranes consisted of DMPC and DMPS, representative of phospholipid classes located in the outer monolayers of cell membranes^*12, 27*^. We understand that the use of different chain lengths of the unsaturated and saturated lipids is not ideal, and the same chain length should be used for better comparison^*28*^. Therefore, we used DMPS lipid for studying the effect of negative lipid component. Our work showed a noble mechanistic pathway of Sup35 aggregation which is greatly modulated by the lipid concentration and composition.

Our results suggest that at a particular lipid/protein ratio (defined as OLC, Optimum Lipid Concentration from now onwards) the aggregation kinetics followed an interesting biphasic pattern. In contrast, the aggregation profile of Sup35 attained the monophasic nature when the aggregation occurred in the presence of high (ELC, Excess Lipid Concentration) and low lipid/protein ratios (LLC, Low Lipid Concentration). The kinetics is monophasic also in the absence of lipid. The presence of negatively charged membrane also showed monophasic aggregation kinetics of Sup35.

## Results

### The model structure of Sup35NM

For the present study, we used the NM region of Sup35NM protein, which has been shown by in vivo and in vitro studies to govern the prion status of the protein ^*19, 29-31*^.

This NM domain is responsible for the formation of toxic amyloid structures after a lag phase^*19*^. Since the understanding of the lipid binding site would require structural insights of the protein, we developed a model structure using its sequence homology with structurally known prion proteins including that from *S. pompe* (Figure S1A) and human (Figure S1B). For the model generation, we used a composite approach through iterative threading assembly refinement (ITASSER) of the Zhang lab server. This method combined threading, ab-initio modelling, and atomic level structure refinement methods, and the model with the best confidence score was accepted as the optimum structure (Figure 1).

**Figure 1.**
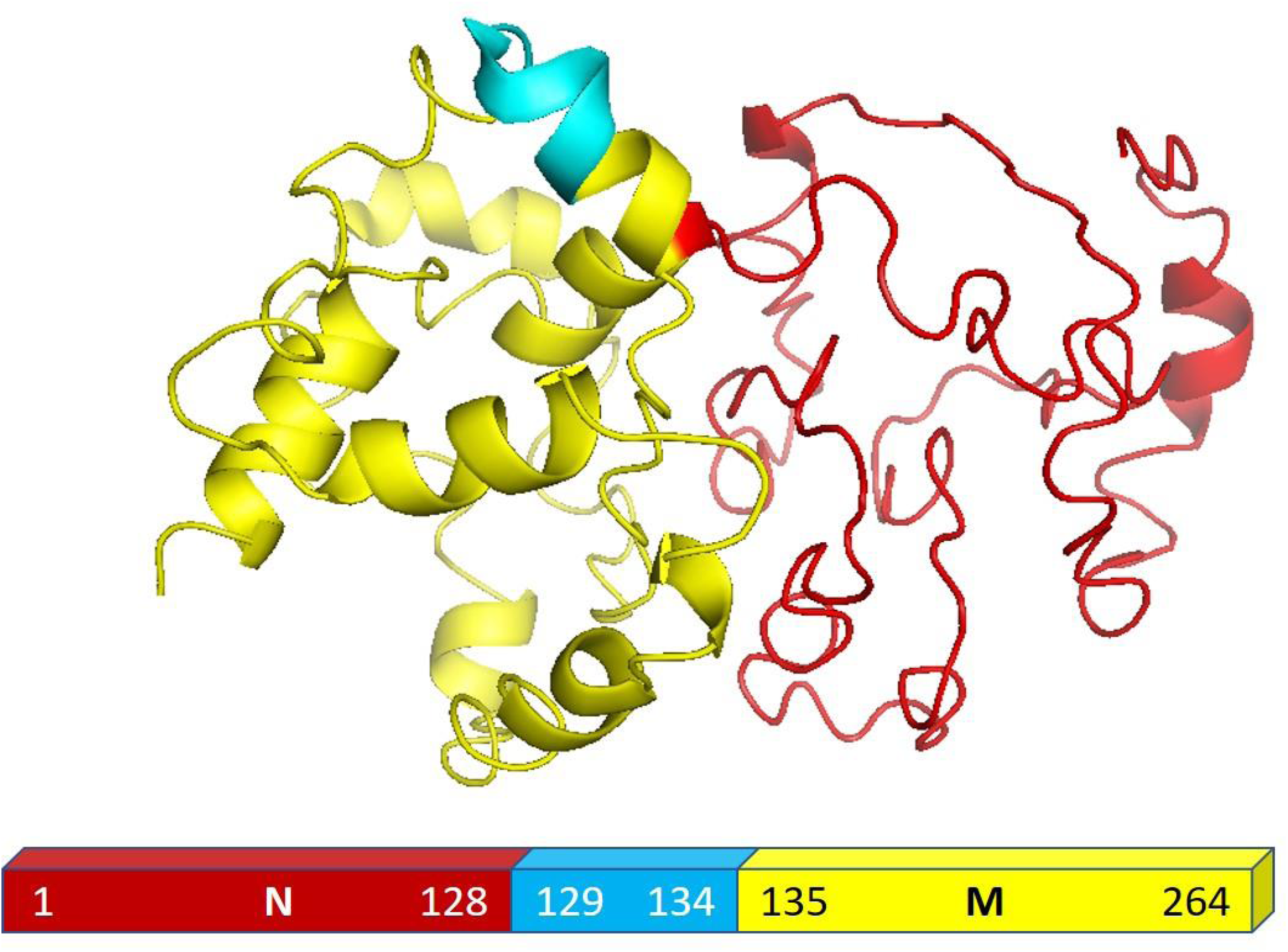
Structural modelling of Sup35NM. This model structure provided the best confidence score from the I-TASSER analyses. The structure of NM region of Sup35NM has been given. The palindromic sequence stretch^129^[QKQQKQ]^134^ in the NM region of Sup35NM has been marked blue.

We validated the model structure using a number of computational methods. Ramachandran plot analysis (https://swift.cmbi.umcn.nl/cgi//PictureCGI.py) showed that all residues in the model occupied the generously allowed region. In the generously allowed region, 70 % of the residues were found inside the favored region (Figure S1C), while the rests were predominantly turns, loops and disordered. In addition to the Ramachandran analyses, we also used proSA algorithm (https://prosa.services.came.sbg.ac.at/prosa.php) to study the overall quality of the model (Figure S1D), using which we compared the Z score of the present model against all other experimentally determined structures available in current PDB database. Z score represents the deviation of total energy of the structure with respect to an energy distribution derived from random conformations. The NMR and X-ray crystal structures in Figure S1D have been shown using dark and light blue colors respectively, while the black dot represents the predicted model of Sup35NM. We observed that the Z score value of the predicted model was located well within the region of structures determined by X ray methods. The plot of the local model quality predicted by proSA program by plotting energies Vs amino acid sequence over the window of 40 residues and 10 residues showed that the energy values of all residues in our predicted model are under zero (Figure S1E).We estimated that the model structure would contain approximately 25% alpha helix and 2% beta sheet while the rest would be random coil/disordered. It has been previously shown that the short prion recognition elements within the N-terminal domain (N), termed as the Head and the Tail may form homotypic intermolecular contacts. While both regions undergo nucleation, the Head region is known to acquire the productive interactions first^*32*^.

### Interaction of Sup35NM with lipid vesicles

To obtain a preliminary understanding of the membrane binding affinity of Sup35NM, we used a computational method using OPM (Orientation of protein in membrane) server. Using the model structure as its input, OPM server calculated the free energy of membrane binding to be −3.6 Kcal/mole (Figure 2A). The calculation predicted that the sequence stretch between 130 and 143, which is mostly a flexible loop region, would potentially be the membrane interacting region. We observed an average tilt angle of 53° between the membrane surface and the interacting stretch of the protein (Figure 2A). Interestingly, we found in this sequence stretch of Sup35NM the presence of a polar palindromic sequence stretch^129^[QKQQKQ]^134^, which may have an important role to play. We noted that a hydrophobic palindromic sequence stretch^112^[AGAAAAGA]^119^ in the human prion protein is found to be a key motif for membrane binding and aggregation progression.

**Figure 2.**
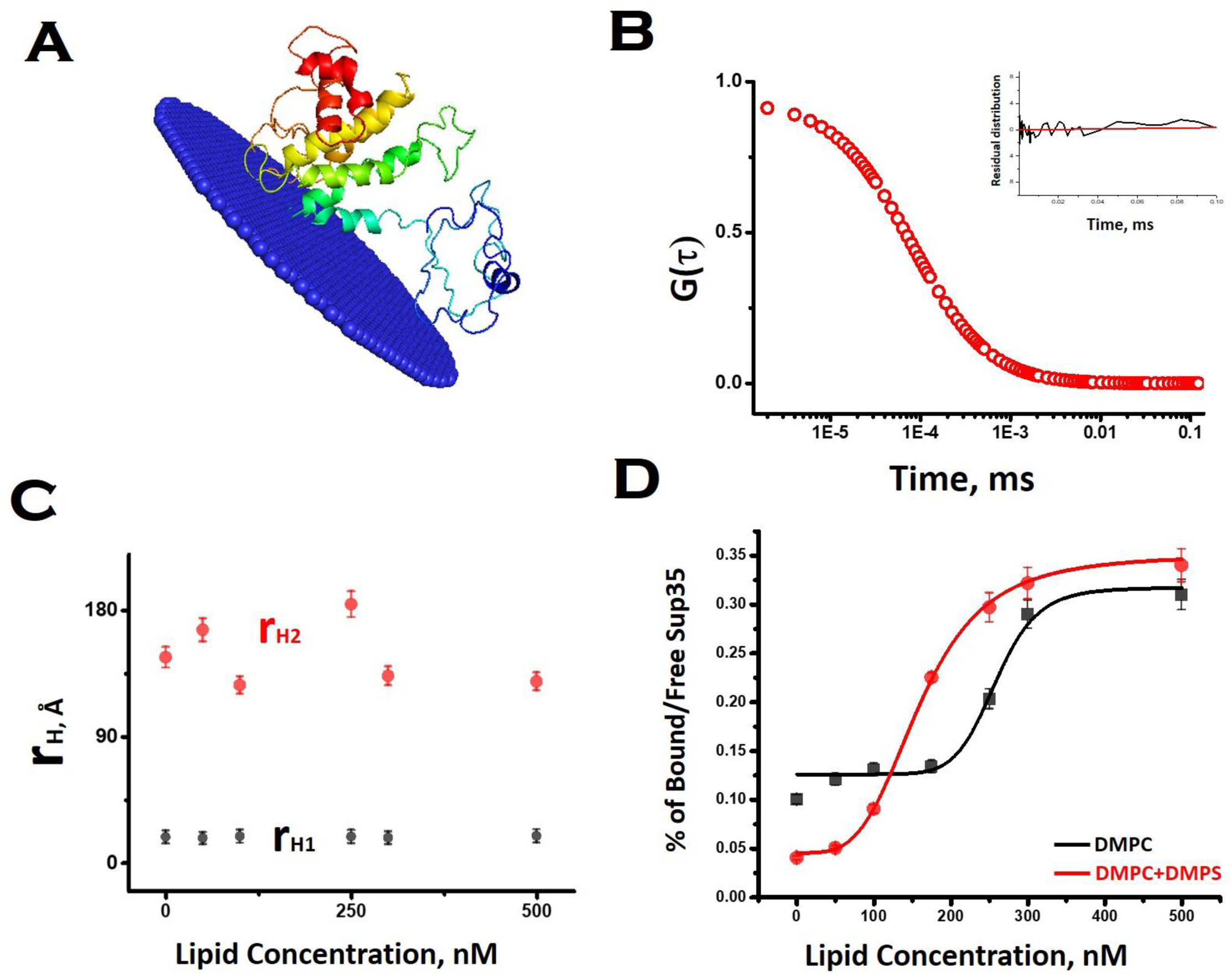
Interaction of Sup35NM with lipid vesicles. **(A)** OPM (Orientation of protein in membrane) server to calculate the free energy of membrane binding of Sup35NM. **(B)** Correlation function obtained from FCS experiments using 10nM Rho-Sup35NM in aqueous buffer at pH 7.5. Inset shows the residual distribution with respect to time. **(C)** r_H_ values for both components which remained constant throughout the concentration range of DMPC SUVs. **(D)** The increase in the fraction/ratio of the membrane bound form of Sup35 with increasing concentration of lipid, black line for protein binding to DMPC vesicles only and red line for protein binding to phospholipid vesicles composed of DMPC and DMPS (50:1).

To investigate experimentally the membrane binding, we resorted to fluorescence correlation spectroscopy (FCS). For FCS experiments, we labeled the N-terminal of Sup35NM using Rhodamine Green (Rho-Sup35NM). Figure 2B show a typical correlation function, which was obtained from FCS experiments using 10 nM Rho-Sup35NM in aqueous buffer at pH 7.5. In the absence of lipid, the correlation function was fit to a model containing a single diffusion component, and the goodness of the fit was established using the randomness of the residual distribution. Using the value of diffusion coefficient (D) obtained from FCS, we determined the experimental value of r_H_ (r_H, exp_) for Rho-Sup35NM to be 18.5 Å. We compared the value of r_H, exp_ with the theoretical value of r_H_ (r_H,theo_), which we calculated using Hydropro^*33*^ by inputting the model structure of Sup35NM and observed the hydrodynamic radius of Sup35NM to be 16 Å.

In the presence of zwitterionic (DMPC) and negatively charged (DMPS) SUVs, the correlation functions were fit using a model of two diffusing components in which the rapidly diffusing fast component represented the unbound monomeric protein while the slow component would be the membrane bound protein. In the presence of different concentrations of DMPC SUVs, the values of r_H_ corresponding to the fast (monomeric protein, 18.5 Å) and slow (membrane bound, 140Å) components remained constant (Figure 2C).In contrast, as we increased SUVs concentrations, we observed an increase in the fraction of the membrane bound Sup35NM (slow component) with a decrease in the fraction of the free monomer (fast component) (Figure 2D). The increase in the fraction/ratio of the membrane bound form of Sup35 with increasing concentration of lipid is shown in figure 2d, where black line represents protein binding to DMPC vesicles only and red line represents protein binding to phospholipid vesicles composed of DMPC and DMPS (50:1). We used a sigmoidal fitting function to fit the dependence of these fractions with DMPC concentrations to determine the values of association constant (K_a_∼ 4.9 x 10^3^ M^-1^). It was observed that the presence of negatively charged component (DMPC: DMPS 50:1), significantly increased membrane binding affinity (K_a_= 6.25 x 10^6^M^-1^).

We used far UV CD to investigate the change in protein secondary structure as a result of membrane binding^*34*^. Far UV CD experiments show double minima at 208 nm and 222 nm suggesting alpha helical structure (Figure 3A). In presence of DMPC vesicles, CD data suggested a decrease in helical structure (Figure 3A). In the presence of negatively charged vesicles (DMPS/DMPC 50:1), the helical content of Sup35NM decreased further. Far UV CD data were complemented by measuring FT-IR. From the carbonyl (C=O) stretching vibrations of amide-I (beta sheet: 1633-38 cm^-1^, alpha Helix: 1649-56 cm^-1^, disordered (1644cm^-^1), and turns and loops (1665-1672cm^-1^), we calculated the percentages of secondary structure (Figure 3B).We found from the FT-IR data that Sup35NM contained 35% alpha helical structure, which was similar to the helical contents determined from the model structure (25%). Similar to what was observed with far UV CD, FT-IR analyses suggested a decrease in helical content (to 22%), as DMPC vesicles were added (Figure 3C). The alpha-helical content of Sup35NM was found to be about 19% in the presence of lipid vesicle composed of DMPC/DMPS 50:1 (Figure 3D).

**Figure 3.**
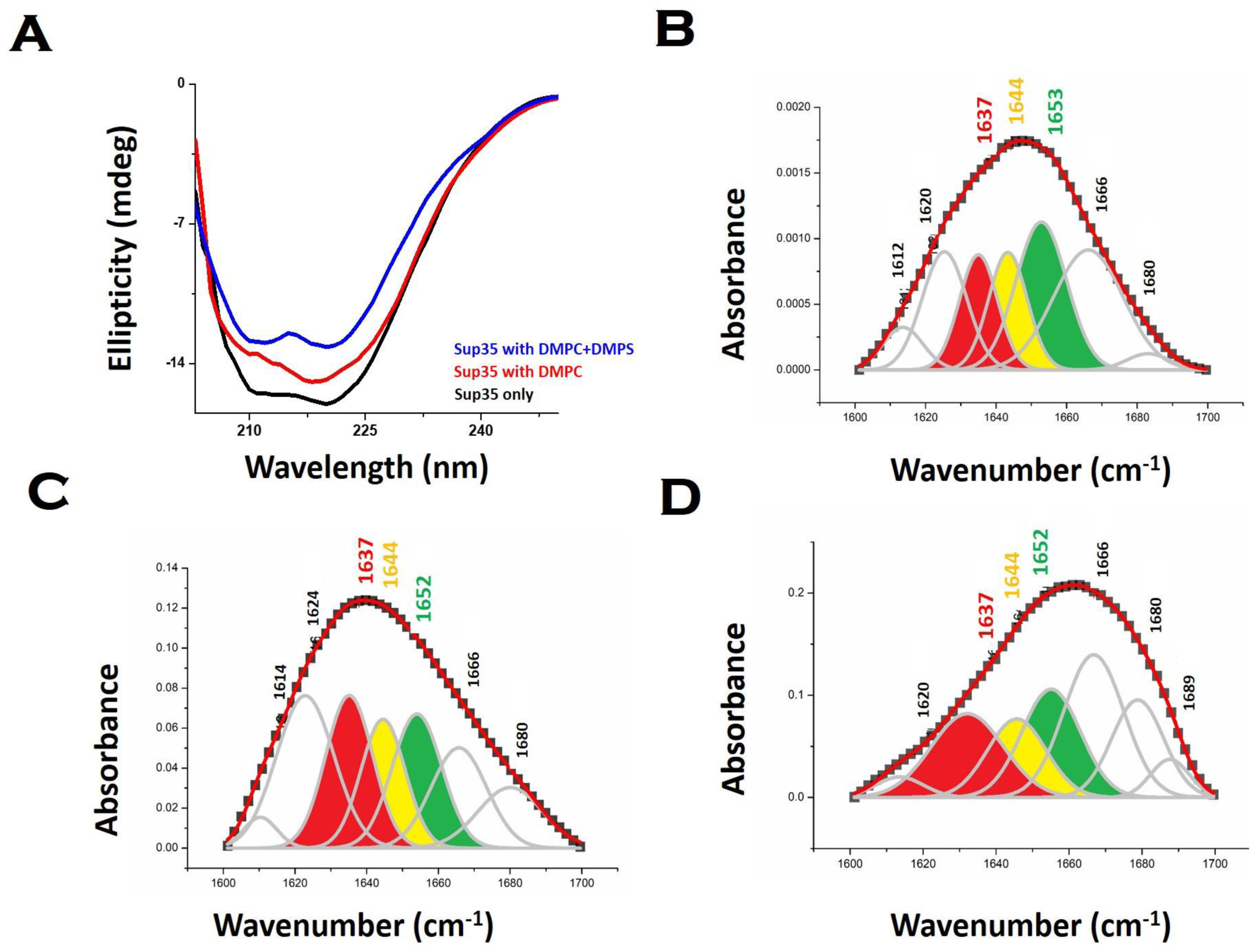
Membrane binding leads to change in Sup35NM structure. **(A)** Far UV-CD spectra of Sup35NM in the absence and presence of lipid. FTIR spectra of the amide-I region of C=O vibrational frequency of Sup35NM in the **(B)** absence of lipid, **(C)** presence of DMPC and **(D)** the presence of phospholipid vesicles composed of DMPC and DMPS (50:1). The color-coded regions signified different conformational states of the protein. Red region indicates the beta sheet conformation; yellow region belongs to the disorder conformation whereas the green region stands for the alpha helical structure.

### Aggregation kinetics of Sup35NM in the absence and presence of membrane

We subsequently studied the effect of the interacting membrane on the aggregation kinetics of Sup35NM. For the initial assessment of the aggregation kinetics, we used the fluorescence intensity enhancement of Thioflavin T (ThT). ThT is known to bind to protein aggregates with cross beta structure giving rise to a large increase in its fluorescence intensity^*35*^. In the absence of membrane, the aggregation kinetics did not show any significant extent of lag phase, suggesting the possibility of the seeded aggregation. Although, ThT fluorescence assay showed that Sup35NM aggregation kinetics reached the saturation phase at ∼54 hours, we continued the experiment for more than 150 hours to observe the saturation phase of aggregation. Glover et al has recently carried out extensive characterization of Sup35NM aggregation kinetics using Congo-red assay with different concentrations of the protein. They found that the aggregation kinetics became faster as the concentration increased. We found that our aggregation kinetics was somewhat slower than the kinetics they observed using similar concentration^*19*^. There are three main differences in the experimental conditions between the present and Glover et al: (a) the probe: while we used ThT fluorescence, Glover et al measured aggregation kinetics using Congo Red (b) the protein stock: we used protein stock freshly prepared for our measurements while Glber et al used diluted protein from stored stocks. In addition, while we used dialyses to remove the small molecules aggregates, Glover et al used column chromatography using a Q-sepharose fast flow column (c) the assay conditions: for the present assay we used 20mM sodium phosphate buffer in contrast to Glover et al, who used potassium phosphate as their buffer of choice. We are currently trying out different experimental conditions to understand the roles of these and other contributing factors towards Sup35NM aggregation.

Since ThT fluorescence has small dependence on lipid concentrations (Figure S4), suitable background correction was used for each time points of aggregation kinetics carried out in the presence of lipids. In the presence of low lipid to protein ratio (LLC condition), we observed a slight increase in the rate of aggregation (the exponential increase started at earlier time point). Interestingly, the extent of aggregation (as measured by the intensity of ThT fluorescence at the saturation phase) remained similar in both conditions (Figure 4A). In the presence of high lipid to protein ratios of DMPC (OLC condition), both the rate and extent of Sup35NM aggregation increased as the concentration of SUVs increased (Figure 4B). More importantly, we found that in these ratios, the aggregation kinetics consisted of two sigmoidal phases. At ELC condition of DMPC, the rate of aggregation was slightly increased with lipid concentration with small change in the extent of aggregation. In these ratios, the aggregation kinetics became mono-phasic again (Figure 4C). Subsequently, we studied the aggregation kinetics in presence of negatively charged lipid membrane (phospholipid vesicles composed of a mixture of DMPC and DMPS in the ratio of 50:1). Surprisingly, it was found that the aggregation kinetics followed monophasic character in the entire range of L/P ratio (Figure 4D).

**Figure 4.**
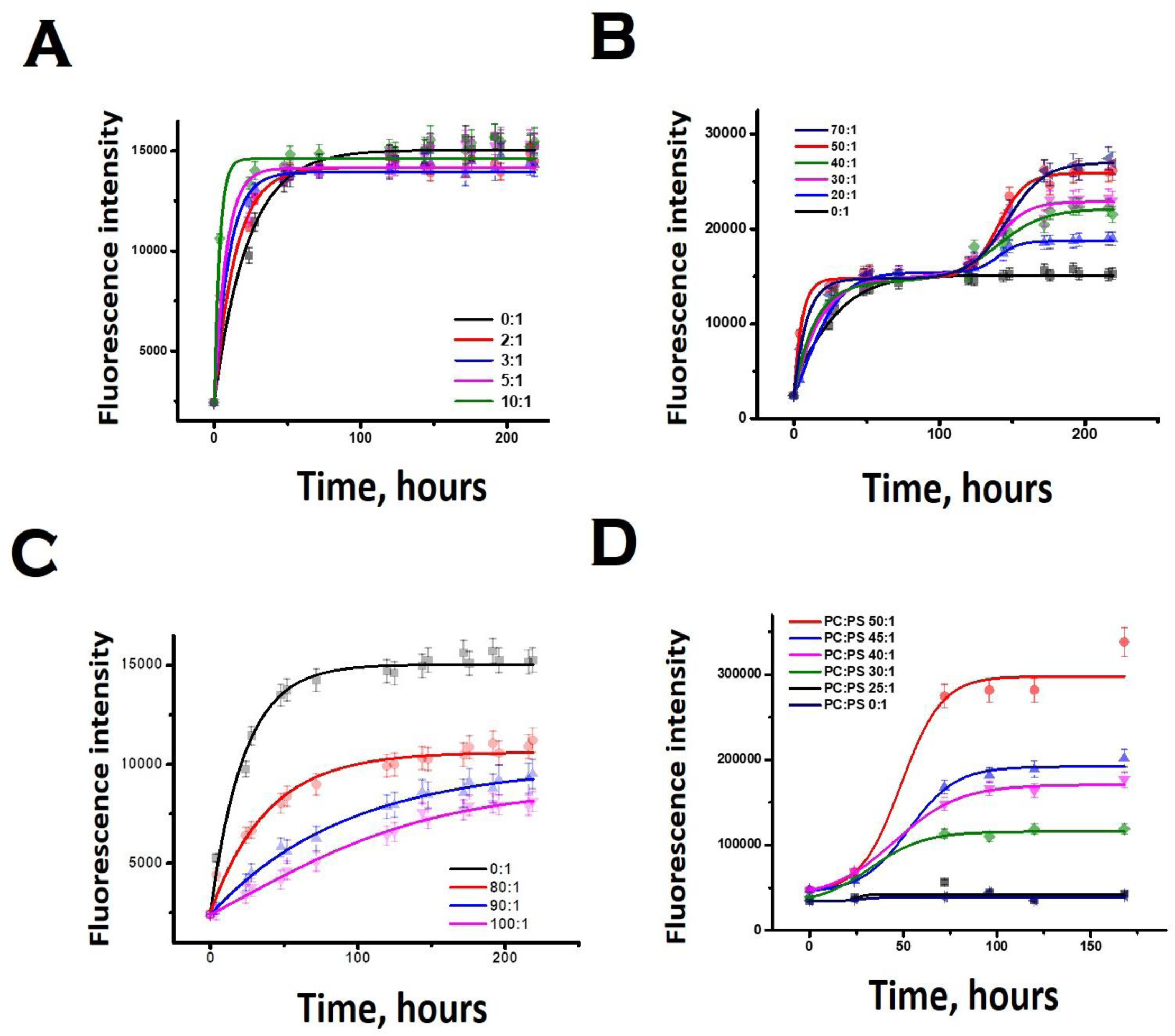
Thioflavin T fluorescence assay showing aggregation of Sup35NM in absence and presence of lipid vesicles. **(A)** At LLC of DMPC. **(B)** Increase in the rate and the extent of Sup35NM aggregation with increase of DMPC concentration at OLC. **(C)** Rate of aggregation was slightly increased with concomitant sharp decrease in the extent of aggregation in the presence of ELC of DMPC. **(D)** Aggregation kinetics in presence of negatively charged lipid membrane (DMPC: DMPS, 50:1 and lower ratios) and by maintaining the L/P ratio at 50:1.

We then complemented ThT aggregation data using FCS. For the FCS experiments, we used 10nM labelled protein in the presence of excess unlabeled Sup35NS for the aggregation to occur. We used three different lipid-to-protein ratios (0:1, 50:1 and 100:1) of DMPC SUVs. Lipid to protein ratio of 0:1 represents the absence of lipid. In the absence of lipids, correlation functions at different incubation time were fit to a sum of two components using the following scheme 1:

### Scheme 1

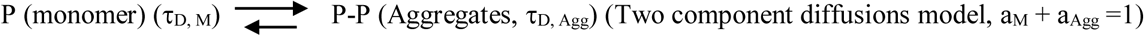

In this scheme, the monomeric protein (denoted by P) would form aggregates (denoted by P-P) and the correlation functions data could be analyzed a two components model with the diffusion time of τ_D,M_ for the monomer (with amplitude of a_M_) and of τ_D,Agg_ (amplitude of a_Agg_) for the aggregates respectively. As expected, a_M_ + a_Agg_=1. Similar terminology and parameter notations would be used subsequently for scheme 2 and 3 below.

Please note that P-P in the above equation indicates a multimolecular aggregates, and not necessarily a dimer. Since FCS data analyses required multiple components and the number of components varied for different conditions, we determined the values of average diffusion times (τ_Dav_) from individual component using Equation 5. Figure 5D show the variations of τ_D,av_ with lipid concentrations for protein in the absence of lipids. The values of τ_D,av_ increased with incubation time following a sigmoidal aggregation kinetics. Comparing the aggregation kinetics monitored by ThT fluorescence and FCS, we noted that they did not superimpose. Although the kinetics measured by both methods saturated approximately at similar hours, ThT monitored kinetics is hyperbolic while FCS monitored one is sigmoidal (Figure S5). In order to get further insights into this difference, we have plotted in Figure 5G, the fractional population of the monomer (P) and aggregates (P-P) with respect to incubation time. As observed in Figure 5G, the amplitude of the aggregated component increased in a sigmoidal fashion, while that of the monomer decreased. The data suggest that the difference (hyperbolic in ThT fluorescence vs sigmoidal in FCS, Figure S5) comes from relatively small fraction of ThT active aggregates at the beginning of the aggregation kinetics. Further studies would be needed to further characterize the nature of these early aggregates.

**Figure 5.**
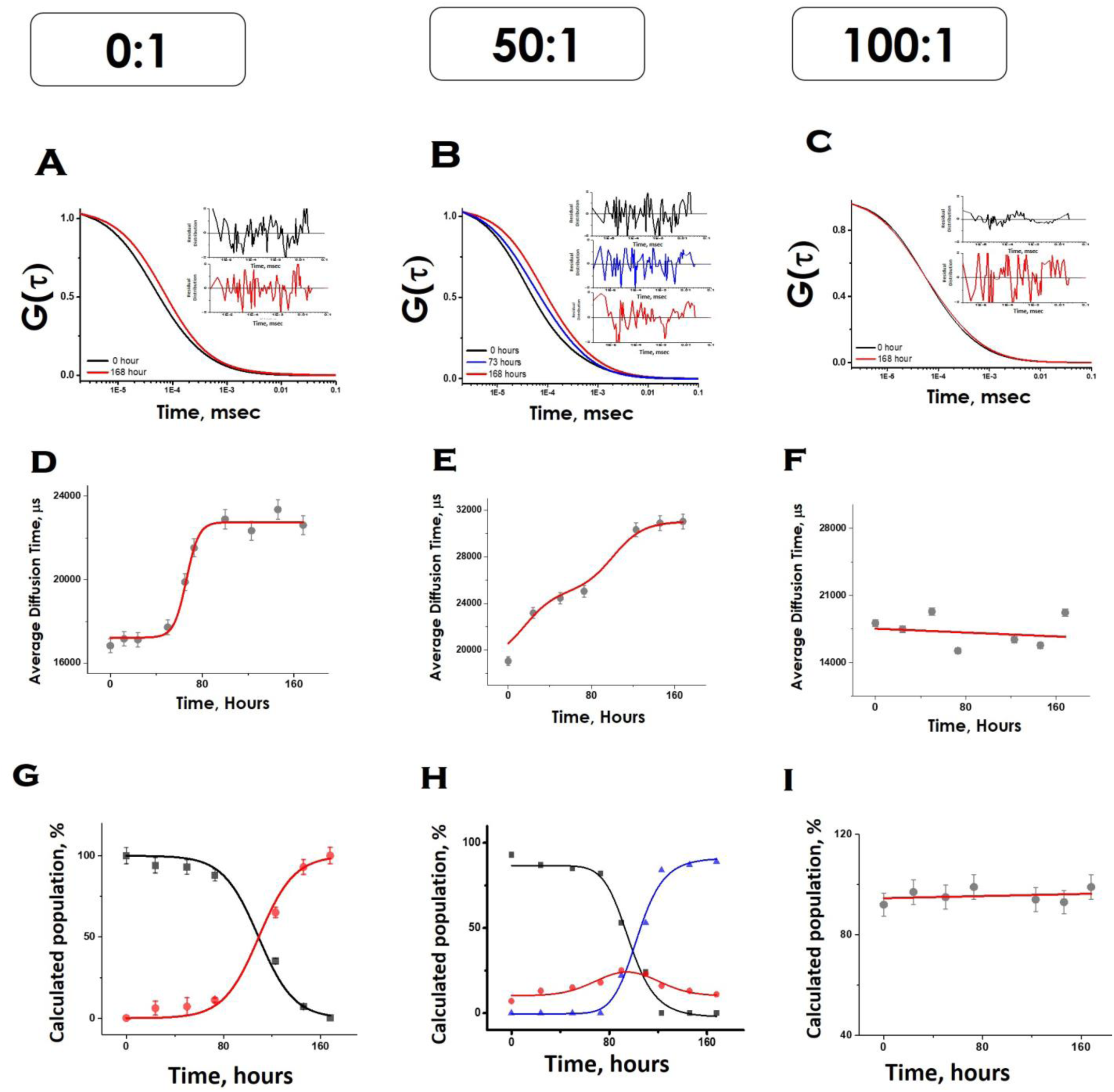
FCS data showing aggregation of Sup35NM in absence and presence of lipid vesicles. The G(τ) was plotted with time as shown in **A, B and C**. The average diffusion time of the protein aggregates were plotted with time as shown in **D, E and F**. The relative populations or amplitudes of fast (monomer) and slow (aggregates) with respect to time were elucidated from their amplitudes as shown in **G, H and I**. In **G and H** black, red and blue lines shows the populations of protein monomers, protein-protein aggregates and membrane bound protein aggregate respectively.

Under OLC condition of DMPC (lipids to protein ratio of 50:1), the correlation functions were initially fit to a two-component diffusion model. Under this condition, the variation of the amplitude of the aggregated component with incubation time followed a two-phase aggregation kinetics (Figure 5E), the trend of which was similar to that observed with ThT fluorescence. We used the following equation scheme (Scheme 2) to analyze the FCS data obtained under this condition:

### Scheme 2

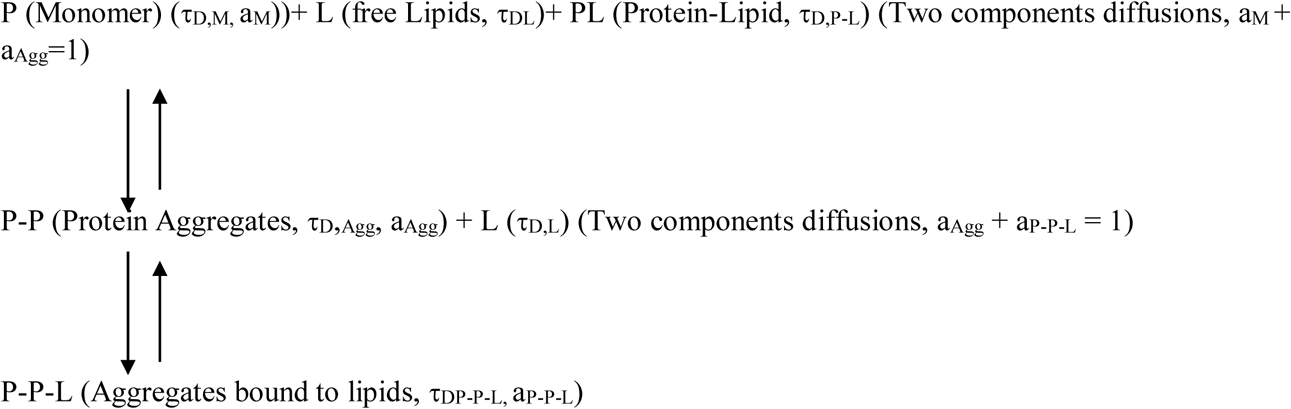

We can ignore τ_D,L_ because lipid is not labeled and hence free lipid would not contribute to the correlation function. In this condition (OLC, the protein remained mostly free at zero time point and at room temperature, and hence τ_D,P-L_ can also be ignored. The first part of the scheme 2 assumes the formation of protein-protein aggregates, which binds to lipid in the second part. The component analyses from the FCS fit show the decrease in the amplitude of the monomeric protein as the amplitude of an intermediate aggregate species (P-P) formed. The amplitude of the intermediate decreased with time, as a final aggregated species (P-P-L) started forming (Figure 5H).

In contrast, in the presence of high DMPC concentration (ELC condition of DMPC, lipids to protein concentration of 100:1), we did not see any variation in the amplitude of the aggregated component with incubation time. As a matter of fact, the correlation functions obtained in this condition were fit optimally using a single component, and the amplitude of this component remained unaltered with incubation time (Figure5I). The following scheme 3 was used to analyze the FCS data for this condition:

### Scheme 3

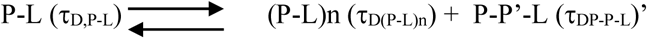

The species (P-L)n indicates the possibility of combining of several protein-lipid complex, which do not seem to be happening The species P-P’-L is also an aggregated protein species, which is bound to lipid molecules (see Scheme 2 above). Since FCS data is fit to only one component diffusion model, the analyses suggest that the species P-L (monomeric protein bound to lipid) and P-P’-L (a protein aggregate bound to the lipid) have similar values of r_H_. This condition is satisfied when a fibril like aggregate is wrapped about a large lipid molecule, in which condition there would not be much difference between a lipid bound monomer and a lipid bound aggregate. In the next section, we have used AFM to validate this possibility.

We used AFM to substantiate the above FCS analyses in the absence and presence of membranes^*36*^. Control DMPC SUVs showed round shaped morphology of diameter ∼ 180 nm (Figure 6A and L in scheme 2 above). In the absence of the membrane, Sup35NM (P in scheme 1) formed aggregates of elongated fibrillar shape (Figure 6B and P-P in scheme 1). The average length of Sup35NM fibrils was found to be ∼2.5µm with a height of 3.4 nm (Figure 6B). In the presence of DMPC at OLP condition, AFM images at the first plateau (∼100 hours) (Figure 6C) suggested the co-existence of few fragmented fibrils (P-P) along with intact lipid vesicles (L). AFM images at the second plateau (∼200 hours) clearly show the association of the vesicles onto the fibrillar aggregates forming a “curved-branched” morphology (Figure 6D). Similar morphology of Sup35NM aggregates were observed in the presence of negatively charged vesicles (DMPC: DMPS, 50:1) at the saturation level (∼100 hours) of aggregation (Figure 6E). The average size/length of Sup35NM aggregates was found to be ∼1.8µm in the presence of membrane (15nm in height). At the ELP conditions, tiny fragmented sup35 fibrils associated with lipid vesicles were observed. AFM imaging showed spherical species of free lipid like particles (Figure 6F), which were bound onto protein aggregates.

**Figure 6.**
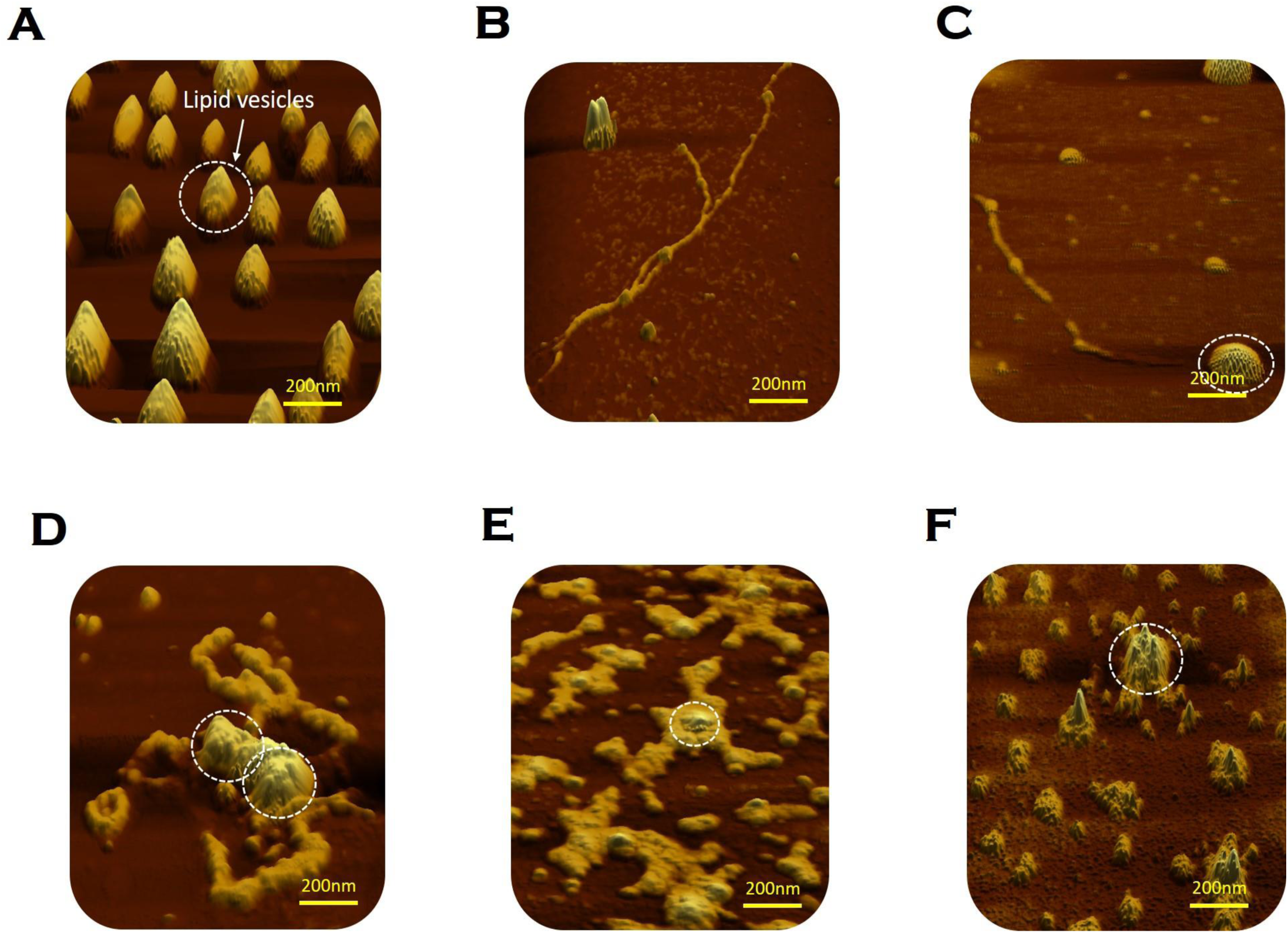
Morphological features of Sup35NM aggregates studied by atomic force microscopy (AFM). **(A)** DMPC SUVs were observed as “round balls”. These lipid vesicles are marked by the white circle. **(B)** Sup35NM aggregates looked like elongated fibrils in the absence of lipid vesicles. **(C)** Co-existence of small Sup35NM fibrils and intact DMPC lipid vesicles imaged at the first plateau (100 hours) of the aggregation curve of lipid to protein ratio of 50:1. **(D)** Curvy and branched fibrillar structures of Sup35NM aggregates were found to protrude around the lipid vesicles when incubated with DMPC SUVs (L/P, 50:1) till 200 hours. **(E)** AFM image of Sup35NM in presence of negatively charged membrane, (DMPC:DMPS, 50:1) by maintaining the L/P ratio at 50, showed curvy and branched fibrillar structures which protruded around the lipid vesicles at 100 hours.**(F)** Existence of a crowd of DMPC lipid vesicles associated with tiny fragmented fibrils were imaged at L/P 100:1 at 200 hours.

### Aggregate-induced toxicity in mammalian cells

Subsequently, we studied the toxicity profiles of Sup35NM aggregates. SH-SY5Y cells were incubated for 12h with Sup35NM aggregates (aggregates formed at 200hrs of incubation). MTT assay was performed under the treatment of Sup35NM aggregates formed in different lipid/protein ratio (0:1, 50:1 and 100:1) of DMPC vesicles. Three different concentrations of each aggregate sample were used for the assay to trace the dose dependent cell viability (Figure S2A). The error bars represent the standard deviation. A slight reduction in the cell viability (89% viability) was observed while the cells were treated with only lipid vesicles. The viable cell population was found to be higher for the treatment of the aggregates in absence of membrane (lipid/protein ratio 0:1), than that of the presence of membrane in the lipid/protein ratio 50:1. In contrast, cell viability was found to be highest after treatment with aggregates, which was formed in the presence of membrane with lipid/protein ratio of 100:1. For each ratio, successive decrease of viable cell population was observed with increase in the concentration of aggregates. Presented results suggested that there was a nice correlation between the cellular toxicity and the lipid/protein ratio. With DMPC L/P ratio of 100:1 being the least toxic and L/P ratio of 50:1 having the highest toxicity toward the mammalian cells.

These experiments were further complemented by calcein release assay using dye entrapped model membrane system. Calcein release assay suggested the fast release and rupture of membrane by the aggregates formed in presence of membrane at the L/P ratio 50:1 (release 83%), (Figure S2B).The rate and extent of dye release was significantly dropped when entrapped vesicles were treated with L/P of 100:1 (release 60%). Whereas Sup35NM aggregates formed in absence of DMPC SUVs showed intermediate release and rupture of membrane (release 72%).

## Discussion

Figure 7 shows the scheme of Sup35NM aggregation in the presence of varying concentration and composition of lipid vesicles. Sup35NM binds both to the membrane (membrane binding regime) and to itself (self-association regime). Lipid to protein ratio of 0:1 is a convenient example of a boundary condition of the self-association regime, in which membrane binding is completely absent. AFM images of Sup35NM aggregates in the absence of membrane looked like elongated fibrils. Next, the conditions with lipid to protein ratio of 50:1 and 100:1 (where membrane was present) were convenient examples, which showed that the membrane binding regime and aggregation regimes modulated each other depending on the lipid to protein ratio. At lipid to protein ratio of 100:1 with DMPC vesicles (ELC condition), where there was an excess amount of lipids, AFM image showed the existence of a crowd of lipid vesicles associated with tiny fragmented fibrils. However, at lipid to protein ratio of 50:1for DMPC vesicles (OLC condition), AFM analysis unveiled that a stretch consisted of oligomers was formed initially, which got elongated on the surface of the DMPC vesicles exhibiting curvy and branched fibrillar structures which were found to protrude around the lipid vesicles. We performed ITC experiments and found that the binding isotherm showed greater binding affinity for protein-protein interactions in comparison to Sup35NM-membrane interactions (Figure S3). Moreover, lipid molecules (at OLC condition) might provide the increased local concentration of protein molecules which in turn facilitated the short chain fibril formation of Sup35NM. Hence, we believe that, at the initial stage of aggregation, protein-protein interaction predominates over protein-membrane interactions producing oligomeric ensembles of protein molecules. With time, these short chain fibrils interacted with the membrane and formed higher order fibrils. As discussed before, FCS analysis at different time points of Sup35NM aggregation in the presence of DMPC vesicles (at lipid to protein ratio of 50:1), show the existence of various Sup35NM populations (Figure 5H). At around 0-hour, high amount of monomeric population (amplitude of the fast component) of Sup35NM was present. With time, monomeric protein gave rise to protein-protein aggregates (amplitude of the second component). During ∼100^th^ hour, monomeric population was minimized with significant increase in the population of protein-protein-aggregate. With further increase of incubation time, these protein-protein aggregates started binding to the membrane surface resulting in the formation of membrane-associated-protein-aggregates (amplitudes of the third component). Thus, at the final point of aggregation, population of membrane-associated-protein-aggregates was dominantly higher than the population of protein-protein aggregate.

**Figure 7.**
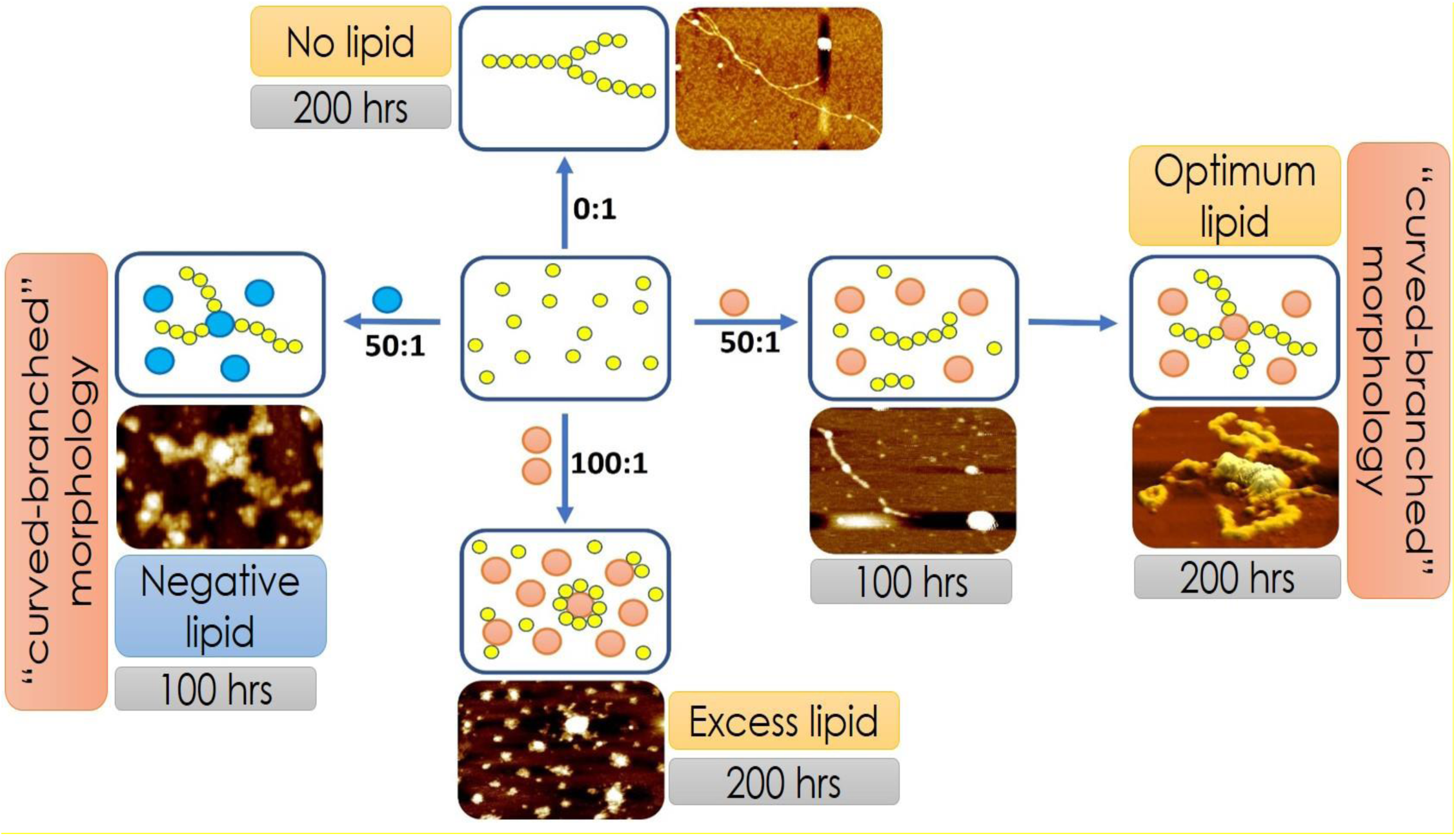
Scheme of Sup35NM aggregation in the presence of varying concentration and composition of lipid vesicles. The membrane binding regime and the aggregation regimes modulated each other depending on the lipid composition and lipid to protein ratio. Yellow circles represent Sup35NM protein molecules. Orange balls represent DMPC lipid vesicles. Blue balls represent lipid vesicles made of a mixture of DMPC and DMPS in the ratio of 50:1.

To obtain a mechanistic insight onto this phenomenon, DMPC vesicles with L/P ratio of 100:1 (ELC condition) was added to the pre-aggregated Sup35NM and incubated for further aggregation which showed a biphasic aggregation profile with ThT-fluorescence assay (Figure S6A). This result further confirmed that already pre-formed Sup35NM aggregates got further elongated on the surface of the vesicles. CD experiments of the pre-aggregated Sup35NM samples were found to exhibit predominantly beta-sheet conformation as confirmed by the existence of major component at 216 nm (Figure S6B). Addition of DMPC vesicles in the L/P ratio of 100:1, significantly altered the secondary conformation from beta sheet to alpha helix. As discussed earlier in the case of alpha synuclein^*37*^, this change in conformation may happen due to dissociation of some Sup35NM protein molecules from the fibrils to populate the lipid-bound monomeric α-helical state.

Nevertheless, Sup35NM aggregation was also significantly modulated by the lipid composition of the membrane. We found that the increase in the negatively charged component (DMPS) significantly increased the aggregation propensity of the protein. It is interesting to note that the biphasic behavior (observed with zwitterionic lipids) disappeared when negatively-charged lipid composition was introduced even at the L/P ratio 50:1. This happened because the presence of anionic lipid component appreciably increased the binding affinity of Sup35NM towards the phospholipid membrane.

We have shown that a good correlation exists between membrane binding affinity, the ability of the aggregates towards pore formation and their extent of cytotoxicity. The extent of membrane deformation and pore formation correlated well with the reduction in cell viability. Aggregates formed in presence of membrane was significantly more toxic in nature than those formed in the absence of membrane. So, our results have provided a mechanistic description of membrane-concentration/composition-modulated-aggregation of Sup35NM and shed some light on aggregation-moderated-cellular-toxicity which are associated with neuronal disease.

## Materials and methods

### Chemicals

DMPC and DMPS were purchased from Avanti Polar Lipids Inc. (Alabaster, Alabama, USA). The amine-reactive compound, Invitrogen Rhodamine Green Carboxylic Acid, Succinimidyl Ester, Hydrochloride (5(6)-CR 110, SE), mixed isomers, Catalog number: R6107 was obtained from Invitrogen (Eugene, USA) and acrylamide, from Sigma-Aldrich (St. Louis, USA). All other necessary chemicals were obtained from Aldrich (St. Louis, USA) and Merck (Mumbai, India).

### Purification of Sup35NM protein

Purification of Sup35NM was performed by following the protocol of Glover et al^*19*^. Recombinant Sup35NM was over-expressed in E. coli (BL21 DE3 strain). The plasmid used for Sup35NM overproduction in E. coli is pJCSUP35NM (1-253) (Addgene Plasmid number #1089). The over-expression of Sup35NM was induced using 1 M IPTG. After induction, the cells were allowed to grow for 3.5 hours. The cells were pelleted down by centrifuging at 6000 rpm for 15 minutes at 4 degrees, which was followed by resuspension in pre-chilled lysis buffer (20 mM Tris-HCl + 500 mM NaCl, pH 8.0). The purification of Sup35NM protein was done in the presence of 8M urea. After thorough re-suspension in lysis buffer the cells were subjected to sonication (20 pulses, each for 30 seconds pulse time and an interim time frame of 1 minute). Unbroken cells and debris were removed by another act of centrifugation at 10,000 rpm for 10 minutes. The soluble fraction obtained thereafter was carefully removed and allowed to bind to Ni-NTA agarose resin. The Ni-NTA column was washed with 40 ml wash buffer (20 mM Tris-HCl, 500 mM NaCl and 50 mM imidazole, pH 8.0) followed by elution with 20 mM Tris HCl, 500 mM NaCl and 500 mM imidazole, pH 8.0. The eluted fractions were pulled according to their tentative protein content as per their absorbance at 280 nm. The post elution fractions were subjected to dialysis in 20 mM Na-phosphate buffer pH 7.5. In all our protein concentration measurements UV-VIS was deployed and Sup35NM concentration was determined by considering the monomeric molar extinction coefficient of 29,000 M−1 cm−1 at 280 nm. The identity of the protein was confirmed by SDS PAGE ^*19*^.

### Preparation of lipid vesicles

An appropriate amount of lipid in chloroform (concentration of stock solution is 25 mg mL^−1^) was transferred to a 10 mL glass bottle ^*26*^. The organic solvent was removed by gently passing dry nitrogen gas. The sample was then placed in a desiccator which is connected to a vacuum pump for couple of hours to remove traces of leftover solvent. Required volume of phosphate buffer saline (PBS) at pH 7.4 was added to the dried lipid film so that the final desired concentration (10 mM) was obtained ^*38*^. The lipid film with the buffer was kept overnight at 4 °C to ensure efficient hydration of the phospholipid heads. Vortexing of the hydrated lipid film for about 30 min produced multilamellar vesicles (MLVs). Long vortexing was occasionally required to make uniform lipid mixtures. Small unilamellar vesicles (SUVs) were prepared by extruding the MLV with LiposoFast using an AVESTIN extruder (Ottawa, Canada). The MLV suspensions were extruded through polycarbonate membranes having pore diameters of 80 and 120 nm. This resulted in the formation of SUVs (average diameter ∼200 nm) with well-defined sizes, as measured by dynamic light scattering (DLS). The vesicle solutions were degassed prior to all measurements, as air bubbles introduced into the sample during extrusion may lead to artifacts.

### FT-IR measurements

FTIR spectra of Sup35NM were acquired using a Bruker 600 series FTIR spectrometer. The FT-IR spectral readouts were collected at pH 7.5 immediately after dispensing the proteins in the buffer solution. Buffer baseline was subtracted before taking each spectrum. The deconvolution of raw spectra in the amide I region (1700 cm−1 to 1600 cm−1) was done using least-squares iterative curve fitting to Gaussian/Lorentzian line shapes. The assignment of peaks was done using previously described spectral components associated with different secondary structures. For investigating the morphological changes of bilayer due to interaction of protein variants and preformed aggregates, FT-IR spectroscopy was utilized by using a typical lipid concentration of 2mM ^*38*^. All possible background corrections were done for each and every experiment.

### AFM studies

Aliquots of aggregating samples were withdrawn after prolonged incubation of ∼200 h at 37 °C and were diluted with 10 mM phosphate buffer, pH 7.4. A 5–8 μl aliquot was taken from the diluted sample and deposited on freshly cleaved mica for 10 min. After removing the excess liquid, the aggregates were rinsed with MilliQ water and then dried with a stream of nitrogen. Images were acquired at room temperature using a Bioscope Catalyst AFM (Bruker Corporation, Billerica, MA) with silicon probes. The standard tapping mode was used to image the morphology of aggregates ^*38*^. The nominal spring constant of the cantilever was kept at 20–80 N/m. The spring constant was calibrated by a thermal tuning method. A standard scan rate of 0.5 Hz with 512 samples per line was used for the imaging of the samples. A single third order flattening of height images with a low pass filter was done followed by section analysis to determine the dimensions of aggregates.

### ThT fluorescence assay

The protein Sup35NM was subjected to mechanical agitation at 200 rpm at 37^0^ C for ∼ 200 hours^*38*^. The protein concentrations for the aggregate preparation were kept 30 µM in 20 mM sodium phosphate buffer at pH 7.5. For the measurement of membrane induced protein aggregation, Sup35NM was incubated at varying protein to lipid ratios from 1:0 to 1:100. Aliquots were thereafter subjected to ThT addition and fluorescence measurements were taken using an integration time of 0.3 s. The steady state fluorescence was monitored using an excitation wavelength of450 nm, and the values of emission intensity at 485 nm were recorded.

### Circular dichroism

Far-UV CD spectra of Sup35NM were recorded using a JASCO J720 spectropolarimeter (Japan Spectroscopic Ltd.). Far-UV CD measurements (between 200 and 250 nm) were performed using a cuvette of 1 mm path length. A protein concentration of 10 μM was used for CD measurements ^*38*^. The scan speed was 50 nm min^−1^, with a response time of 2 s. The bandwidth was set at 1 nm. Three CD spectra were recorded in the continuous mode and averaged.

### Assay for permeabilization of lipid vesicles

The ability of proteins aggregates to cause release of calcein from entrapped DMPC vesicles was checked by monitoring the increase in fluorescence intensity of calcein. Calcein loaded liposomes were separated from non-encapsulated (free) calcein by gel filtration using a Sephadex G-75 column (Sigma). We used an elution buffer of 10 mM MOPS, 150 Mm NaCl and 5 mM EDTA (pH 7.4), and lipid concentrations were estimated by complexation with ammonium ferro thiocyanate. Fluorescence was measured at room temperature (25°C) by a PTI spectrofluorometer using a cuvette of 1cm path length. The excitation wavelength was 490 nm and emission was set at 520 nm. Excitation and emission slits with a nominal bandpass of 3 and 5 nm were used, respectively. The high concentration (10mM) of the entrapped calcein led to self-quenching of its fluorescence resulting in low fluorescence intensity of the vesicles (I_B_). Release of calcein caused by addition of proteins aggregates led to the dilution of the dye into the medium, which could therefore be monitored by an enhancement of fluorescence intensity (I_F_). This enhancement of fluorescence is a measure of the extent of vesicle permeabilization. The experiments were normalized relative to the total fluorescence intensity (I_T_) corresponding to the total release of calcein after complete disruption of all the vesicles by addition of Triton X-100 (2% v/v). The percentage of calcein release in the presence of melittin was calculated using the equation:

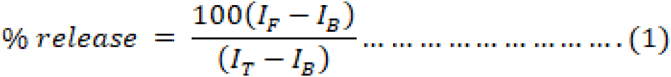

where, I_B_ is the background (self-quenched) intensity of calcein encapsulated in vesicles, I_F_ represents the enhanced fluorescence intensity resulting from the dilution of dye in the medium caused by protein-aggregates induced release of entrapped calcein. I_T_ is the total fluorescence intensity after complete permeabilization is achieved upon addition of Triton X-100^*39*^.

### Cell culture and cytotoxicity assays

SHSY5Y cells were cultured in DMEM accompanied with 10% fetal bovine serum (FBS) and 1% antibiotic (PSN) under 5% CO_2_ in a humidified atmosphere at 37°C^*21, 40*^. After 75–80% confluence, cells were harvested with 0.52 mM EDTA and 0.25% trypsin in phosphate buffered saline (PBS) and plated at the required density to allow them to re-equilibrate for a day before starting the experiment. MTT assay was directed to evaluate the cell cytotoxicity. For the initial screening experiment, the SHSY5Y cells (4×10^3^ cells per well) were seeded in a 96 well plate and left in an incubator followed by treatment with different concentrations of protein aggregates (5,10 and 15µM) for 12 h. After 12 h of incubation, cells were washed with PBS, and then the MTT solution was added to each well and kept in an incubator for 4 h to form formazan salt. Then the formazan salt was solubilized using DMSO and the absorbance was observed at 595 nm using an ELISA reader (Emax, Molecular device, USA).

### OPM (Orientations of proteins in membranes)

To obtain an insight into how the Sup35NM protein interacts with the membrane, we resorted to computational approaches. Protein orientations in membranes were theoretically calculated by minimizing a protein’s transfer energy from water to a planar slab that serves as a crude approximation of the membrane hydrocarbon core. The membrane binding propensity was calculated using OPM server (https://opm.phar.umich.edu/server.php). A protein was considered as a rigid body that freely floats in the planar hydrocarbon core of a lipid bilayer. Accessible surface area is calculated using the subroutine SOLVA from NACCESS with radii of Chothia and without hydrogen. In OPM, solvation parameters are derived specifically for lipid bilayers and normalized by the effective concentration of water, which changes gradually along the bilayer normal in a relatively narrow region between the lipid head group regions and the hydrocarbon core.

### ITC (Isothermal Titration Calorimetry)

The heat flow obtained from the binding of Sup35NM with phospholipid vesicles was measured using high-sensitivity ITC (MicroCaliTC 200, U.K.). All experiments were performed at 30 °C. Solutions were degassed under vacuum prior to filling the sample and reference cells. Protein solution (10 μM) and DMPC SUVs (2 mM) were loaded in a calorimetric cell of volume 350 μL and a syringe of volume 40 μL, respectively. In an experiment, a series of 20 injections, 2 μL each, with the syringe was performed into the ITC cell containing Sup35NM at 240 s intervals. Each injection produced a characteristic heat signal, arising from the released or absorbed heat from the lipid−protein interaction, leading to exothermic and endothermic signals. The heat of dilution was determined by injecting the buffer into the ITC cell containing Sup35NM. An ITC thermogram was obtained from the integration of a heat signal and subtracting the heat of dilution arising from each injection. The ITC thermogram was then fitted to a model (one-site binding) provided by Microcal Origin to determine the binding constant (K) and molar enthalpy of interaction (ΔH). The Gibbs free energy, G, was obtained from the relation ΔG = −RTln(55.5K) and hence the entropic contribution can be estimated as follows: ΔG = ΔH − TΔS. Here, the concentration of water of 55.5 M was used to correct the unit of K to molar fraction.

### Labelling of SUP35NM

10 mg of the protein was dissolved in 1 mL of 0.1 M sodium bicarbonate buffer. The protein concentration in the reaction was 10 mg/mL. The amine-reactive compound, Invitrogen Rhodamine Green Carboxylic Acid, Succinimidyl Ester, Hydrochloride (5(6)-CR 110, SE), mixed isomers, Catalog number: R6107 (Rhodamine dye), was dissolved in DMSO at 10 mg/mL.Vortexing and sonication was done. Incubation of the reaction was done for 1 hour at room temperature with continuous stirring. Equilibration was done of a 10 × 300 mm column with PBS or buffer of choice. Measurement of the absorbance of the protein–dye conjugate at 280 nm (A280) was taken and at the λmax for the dye.

### Fluorescence Correlation Spectroscopy

FCS experiments were carried out using a dual channel ISS Alba V system (ISS, IL) equipped with a 60× water-immersion objective (NA 1.2). Samples were excited with an argon laser at 488 nm. All protein data were normalized using the τ_D_ value obtained with the free dye (Alexa488) which was measured under identical conditions. For a single-component system, diffusion time (τ_D_) of a fluorophore and the average number of particles (N) in the observation volume can be calculated by fitting the correlationfunction [G(τ)] to Eq. 2:

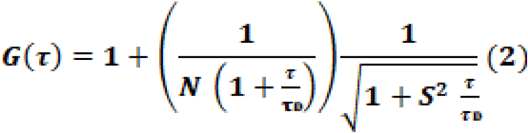

where, S is the structure parameter, which is the depth-to-diameter ratio. The characteristic diffusion coefficient (D) of the molecule can be calculated from τ_D_ using Eq. 3:

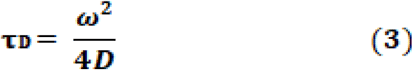

where, ω is the radius of the observation volume, which can be obtained by measuring the τ_D_ of a fluorophore with known D value. The value of hydrodynamic radius (r_H_) of a labelled molecule can be calculated from D using the Stokes–Einstein equation [Eq. 4]:

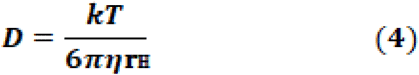

where, k is the Boltzmann constant, T is the temperature and η corresponds to the viscosity of the solution.^*41*^

Calculation of the average diffusion time was carried out using the equation 5, where A_1_ and τ_1_ are the amplitudes and diffusion times of the fast component and A_2_ and τ_2_ are the amplitudes and diffusion times of the slow component.

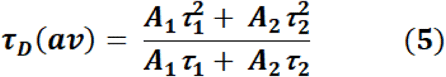

## Supporting Information

Validation of the model structure of Sup35NM, aggregate-induced differential toxicity of Sup35NM in absence and presence of DMPC vesicles, ThT fluorescence showing binding of ThT to lipid vesicles onlyand ITC experiments showing binding affinities, average diffusion time of aggregates superimposed with ThT fluorescence intensity.

## Acknowledgement

This work received funding from CSIR (MIND), Govt. of India. AS and AB are grateful to UGC for providing their research fellowship. The authors thank the Central Instrument Facility of CSIR-IICB for providing the instruments and facilities. AB thanks Mr. T. Muruganandan, technical officer in charge of AFM instrument for helping out with the AFM images. AB thanks Trishika, Basudha, Sanchari (project-trainees) for their help and support. KC thanks the director, CSIR-IICB for his help and encouragements.

## Author contribution

Conceptualization- KC; Supervision- AS; Formal analysis & Investigation- AB; Methodology- AS & AB; Manuscript writing: KC, AS and AB.

## For Table of Contents Only

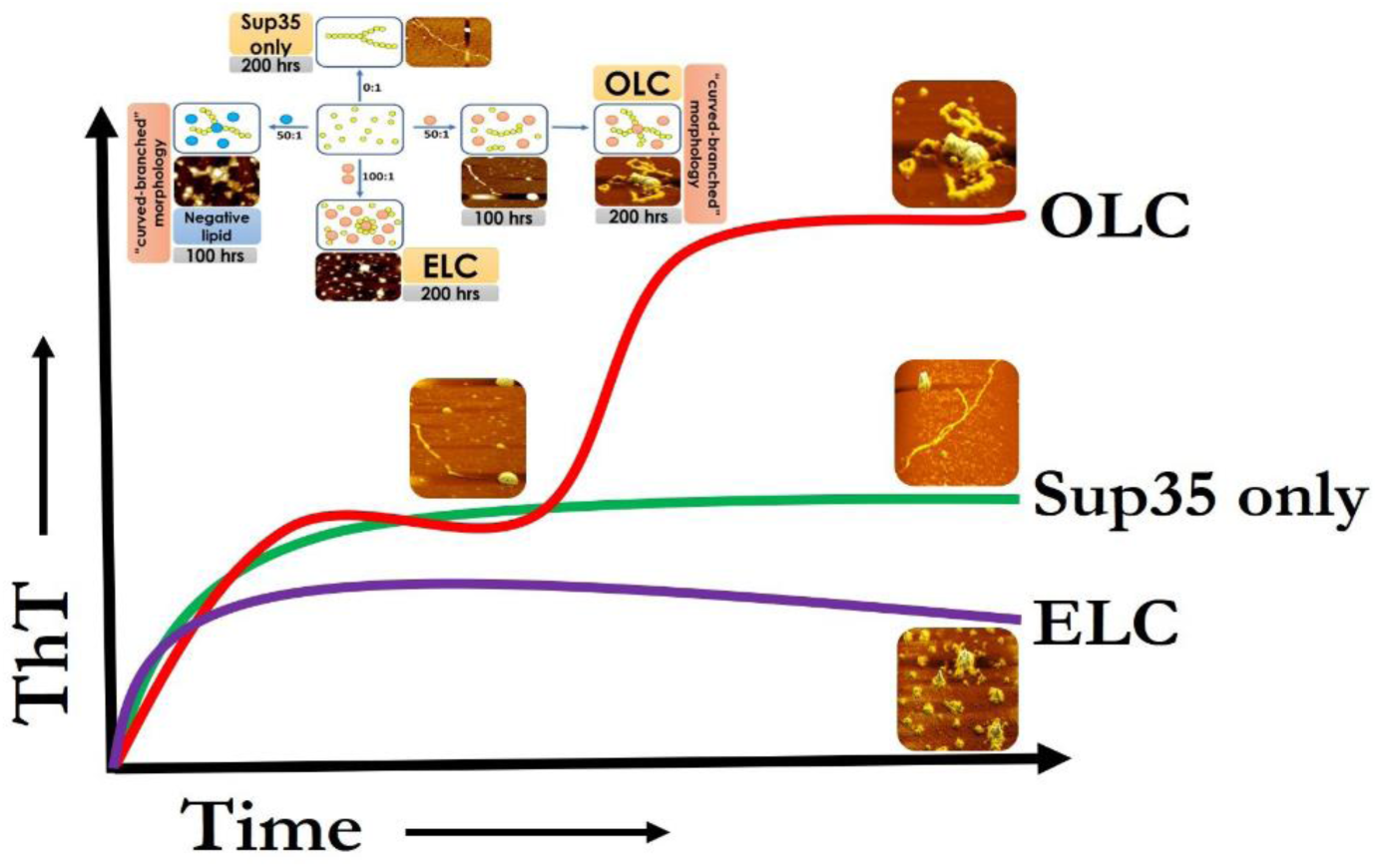

