## Supplemental Figure S1-S6 for "Membrane composition and lipid to protein ratio modulate amyloid kinetics of yeast prion protein"

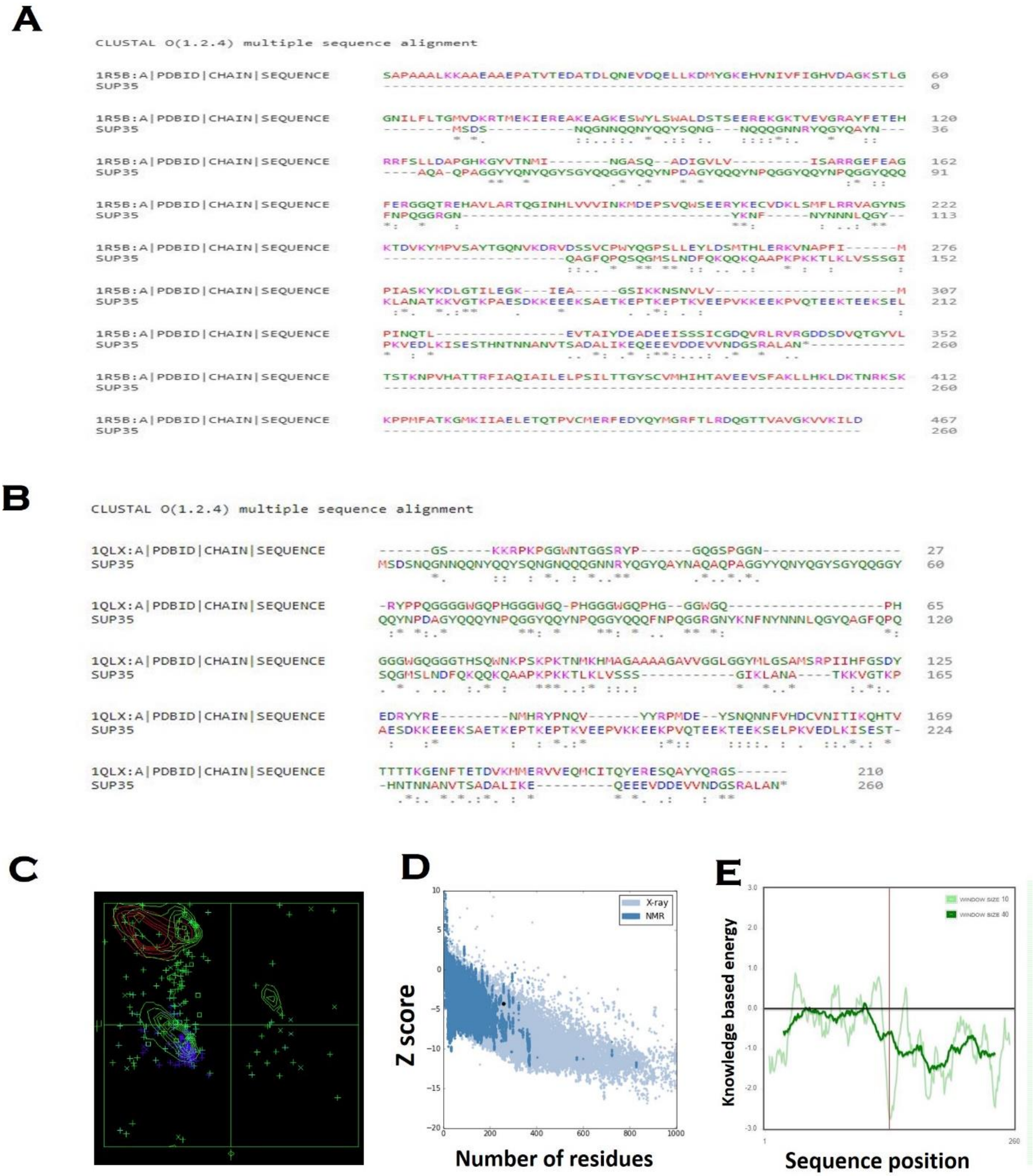


**Figure S1. Validation of the model structure.** Multiple sequence alignment of the sequence of prion protein of S. pompe **(A)** and human prion protein **(B)** with the NM region of yeast prion protein. **(C)** Ramachandran plot analysis of Sup35 protein. **(D)** Z score of our model using proSA algorithm. **(E)** Local model quality predicted by proSA program by plotting energies vs amino acid sequence over the window of 40 residues and 10 residues.

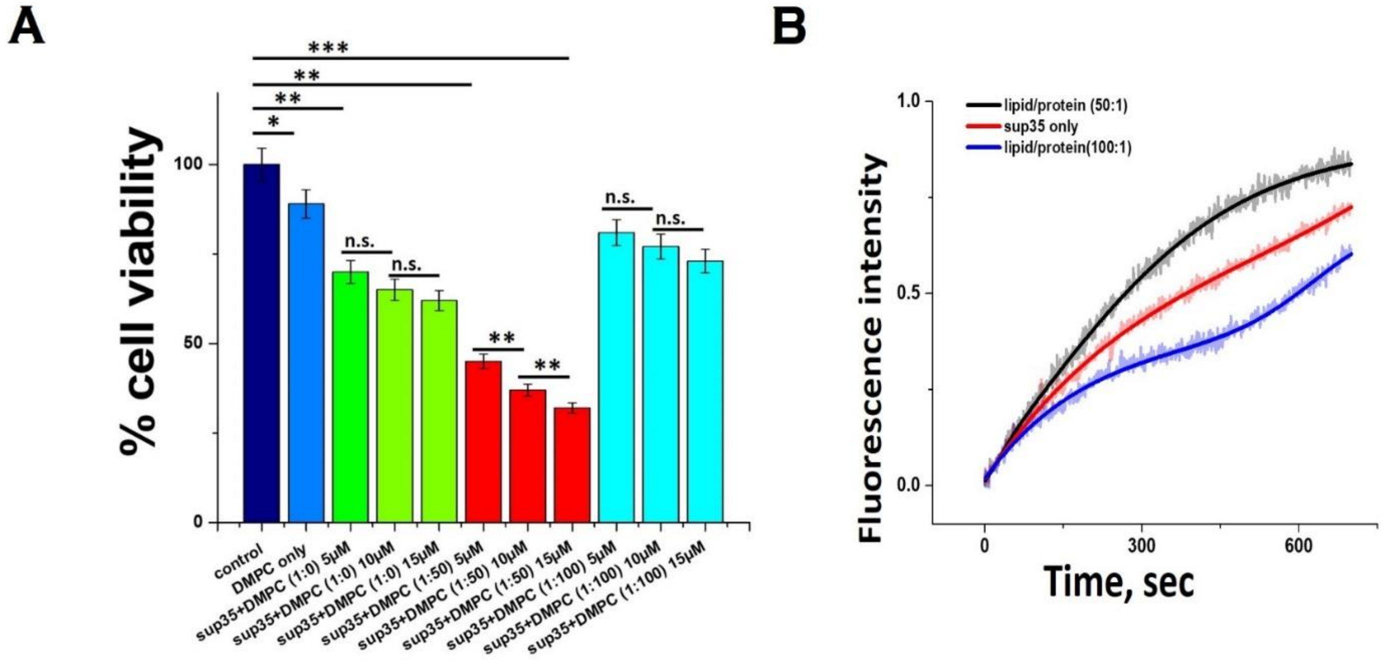


**Figure S2. Aggregate-induced differential toxicity of Sup35 in absence and presence of DMPC vesicles. (A)** 5um, 10um and 15um concentrations of each aggregate sample were used for the MTT assay to trace the dose dependence on cell viability. Error bar indicates the standard deviation. n.s. stands for non-significant data. For the significant changes- *, P value<0.05; **, P value<0.01; ***, P value<0.001. **(B)** Calcein release assay using dye entrapped model membrane system in the presence of lipid /protein ratio of 0:1, 50:1 and 100:1.

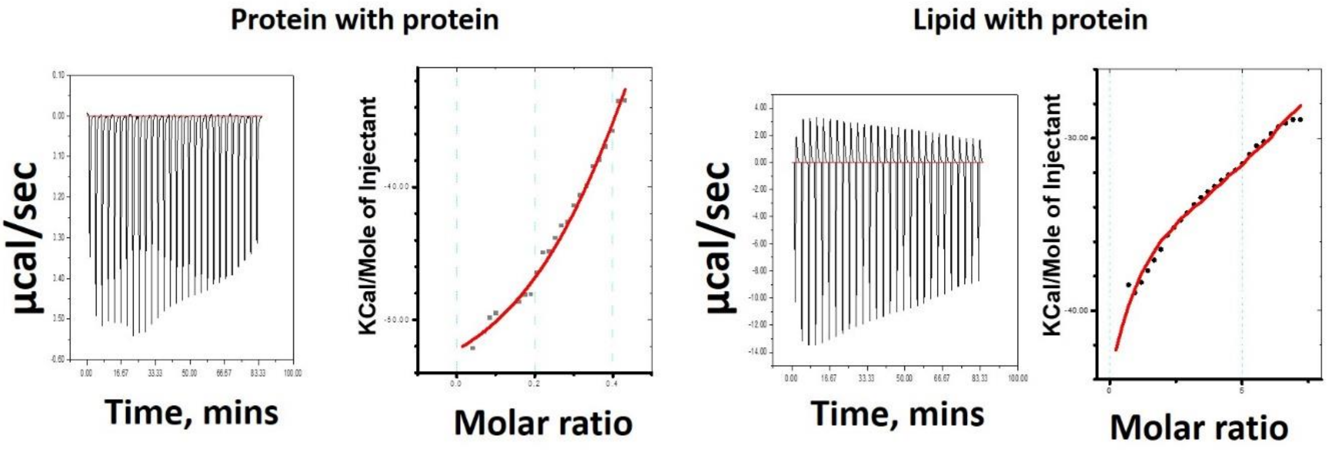


**Figure S3. ITC experiments showing binding affinities.** The binding isotherm showed the greater binding affinity of Sup35 molecules with itself than its affinity towards DMPC vesicles.

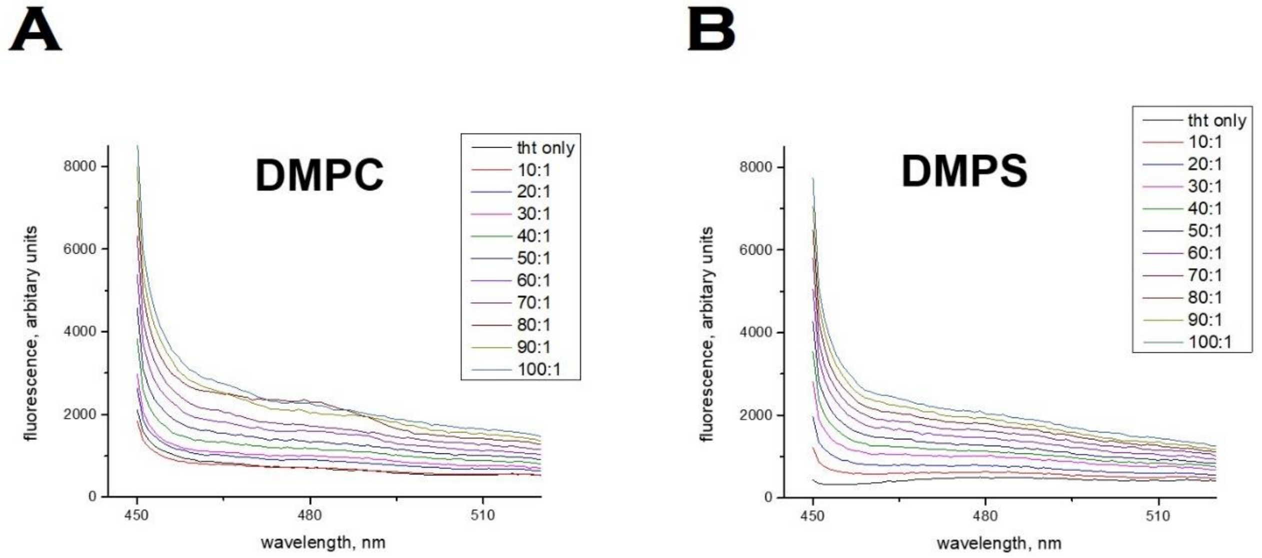


**Figure S4. ThT fluorescence showing binding of ThT to lipid vesicles only. (A)** with neutral vesicles such as DMPC **(B)** with negatively charged vesicles such as DMPS.

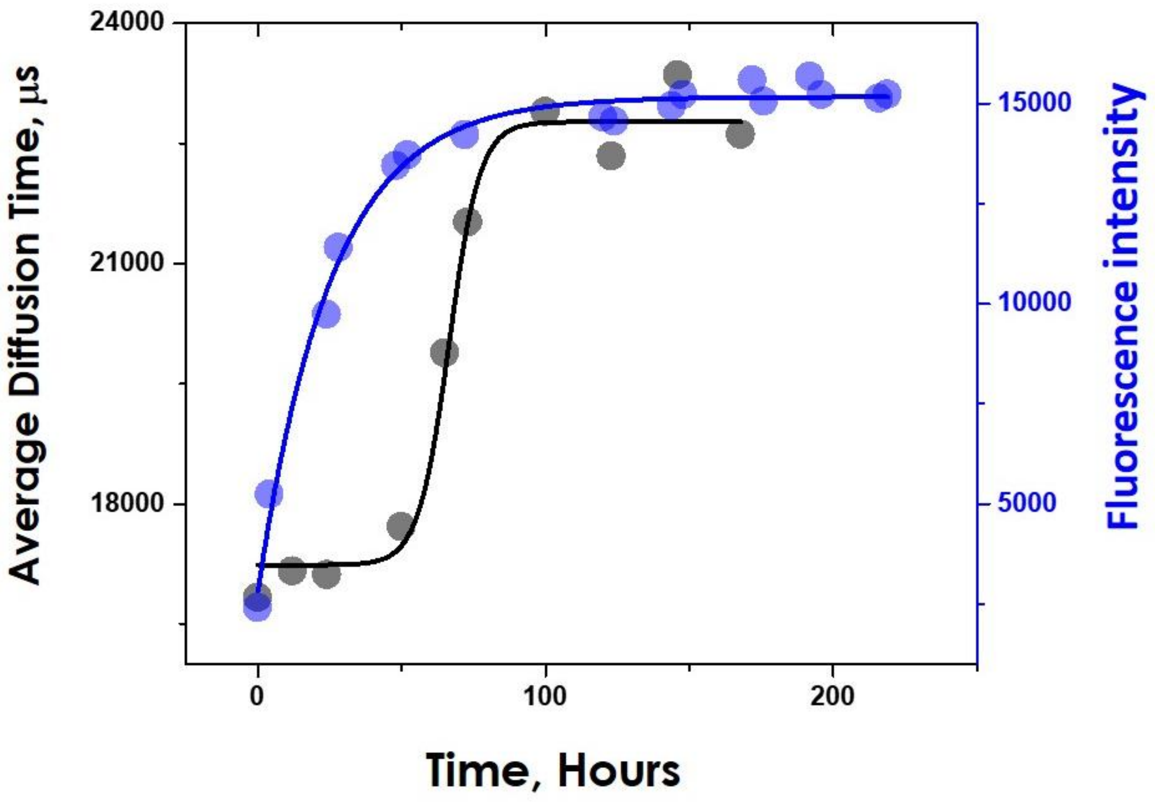


**Figure S5. Average diffusion time of aggregates superimposed with ThT fluorescence intensity.** ThT monitored aggregation kinetics of Sup35 in the absence of lipid is hyperbolic while the FCS monitored one is sigmoidal.

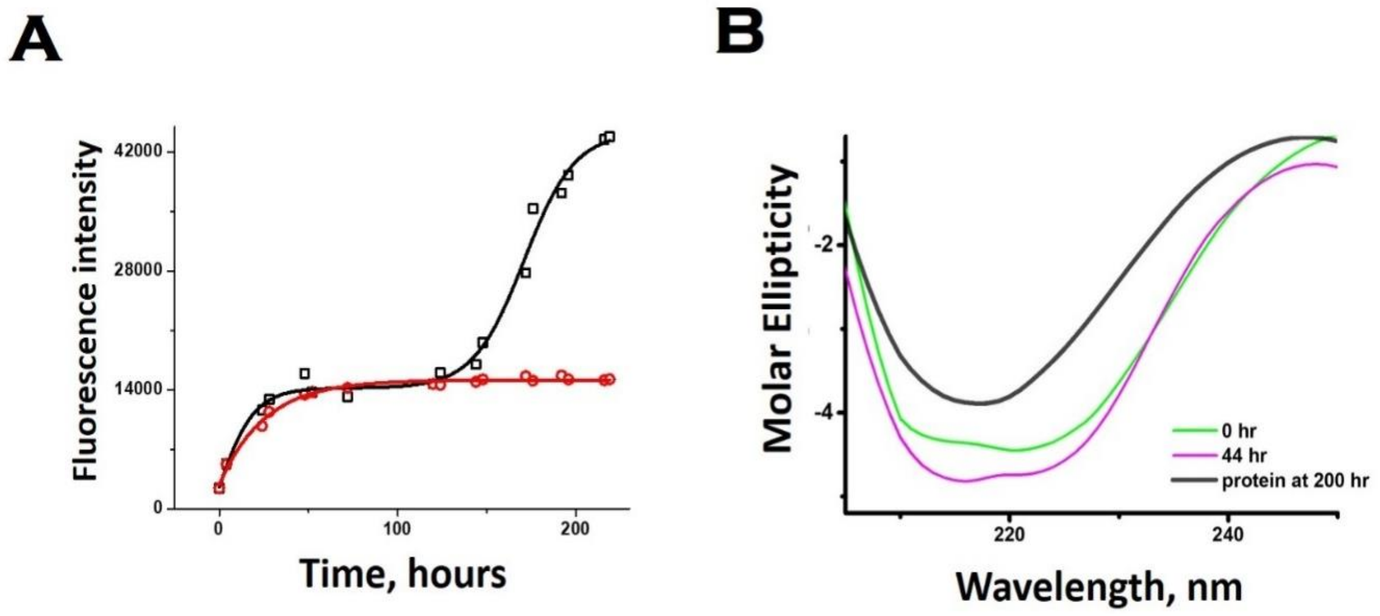


**Figure S6. Mechanistic insight on Sup35NM aggregation. (A)**Assessment of the aggregation using Thioflavin T fluorescence when membrane environment was introduced to the pre-aggregated Sup35NM. Red line represents the aggregation profile of Sup35NM in the absence of membrane. Black line represents the aggregation profile when membrane was introduced at 130 hours. **(B)** Far UV-CD experiments of the pre-aggregated species of Sup35NM when incubated with DMPC SUVs maintaining the L/P molar ratio of 100:1 at different time points of incubation.
